# A horizontally acquired and recurrently expanded glycoside hydrolase subfamily across leafhoppers

**DOI:** 10.64898/2026.07.27.741091

**Authors:** Joshua Molligan, Elisa Ines Fantino, Abraão Almeida Santos, Edel Pérez-Lopez

## Abstract

Horizontal gene transfer from bacteria is a known source of metabolic novelty in insects, yet how these acquisitions diversify and persist over evolutionary time scales remains poorly understood. Here, we reconstructed the evolutionary history of the bacterial glycoside hydrolase subfamily GH5-40 across leafhoppers (Cicadellidae). We annotated 24 genomes and identified 87 GH5-40 genes encoding 113 catalytic domains across 23 leafhopper species, with copy numbers ranging from 1 to 19 genes per genome. Maximum-likelihood phylogenetic analyses recovered all leafhopper GH5-40 domains as a single clade nested within Actinobacteria, supporting one ancestral acquisition followed by extensive lineage-specific duplication of both genes and catalytic domains. Seventeen genes encode 2 to 4 tandem catalytic domains connected by disordered linkers, and a four-domain architecture recurs independently in two divergent leafhopper subfamilies. Recombinant enzymes from distantly related species displayed contrasting substrate preferences for β-glucans and β-mannans *in vitro*, despite GH5-40 enzymes being classically characterized as endo-β-1,4-mannanases.

## INTRODUCTION

Horizontal gene transfer (HGT) from bacteria to eukaryotes has repeatedly contributed to metabolic innovation, with glycoside hydrolase (GH) genes among the best-characterized examples in insects (1,2). Across multiple insect orders, bacteria-derived GH families have been retained and co-opted as digestive enzymes that degrade plant cell-wall polysaccharides, contributing in part to the evolution of herbivory (3–5). Within plant-feeding Hemiptera, however, these GHs remain largely unexplored (6, 7).

The only previous evidence for a GH5-40 enzyme in leafhoppers (Cicadellidae) is the salivary protein NcSP38 from the green rice leafhopper *Nephotettix cincticeps*, which shares close homology with a β-1,4-endoglucanase identified in the glassy-winged sharpshooter *Homalodisca vitripennis* (8, 9). Beyond their identification, neither has been fully characterized or assessed for bacterial origin, and whether they represent independent adaptations or descendants of a shared, more ancient acquisition has remained unclear.

Leafhoppers comprise more than 20,000 described species spanning phloem, xylem, and mesophyll feeders (10–14). With chromosome-level genome assemblies now available across this diversity, they offer an opportunity to trace horizontally acquired gene families across a broad radiation. Here we surveyed GH5-40 across 24 leafhopper genomes, reconstructed the family’s likely origin, and asked whether its expansion has been accompanied by divergence in substrate preference.

## RESULTS AND DISCUSSION

Annotations from 24 leafhopper genomes spanning six Cicadellidae subfamilies recovered GH5-40 in 23 species (**Dataset S1**). In total, 87 genes encoding 113 catalytic domains were identified, with copy numbers ranging from 1 to 19 per genome (**Fig. 1A, Dataset S2**). Eighty-six of the 87 genes had complete structures, with a single locus carrying a premature stop codon, indicating that GH5-40 has been broadly preserved as an intact family across the sampled radiation. Most loci (75/87) were retrieved by standardized *ab initio* annotation, and 12 additional genes from nine species were recovered by protein-guided miniprot search.

**Figure 1.**
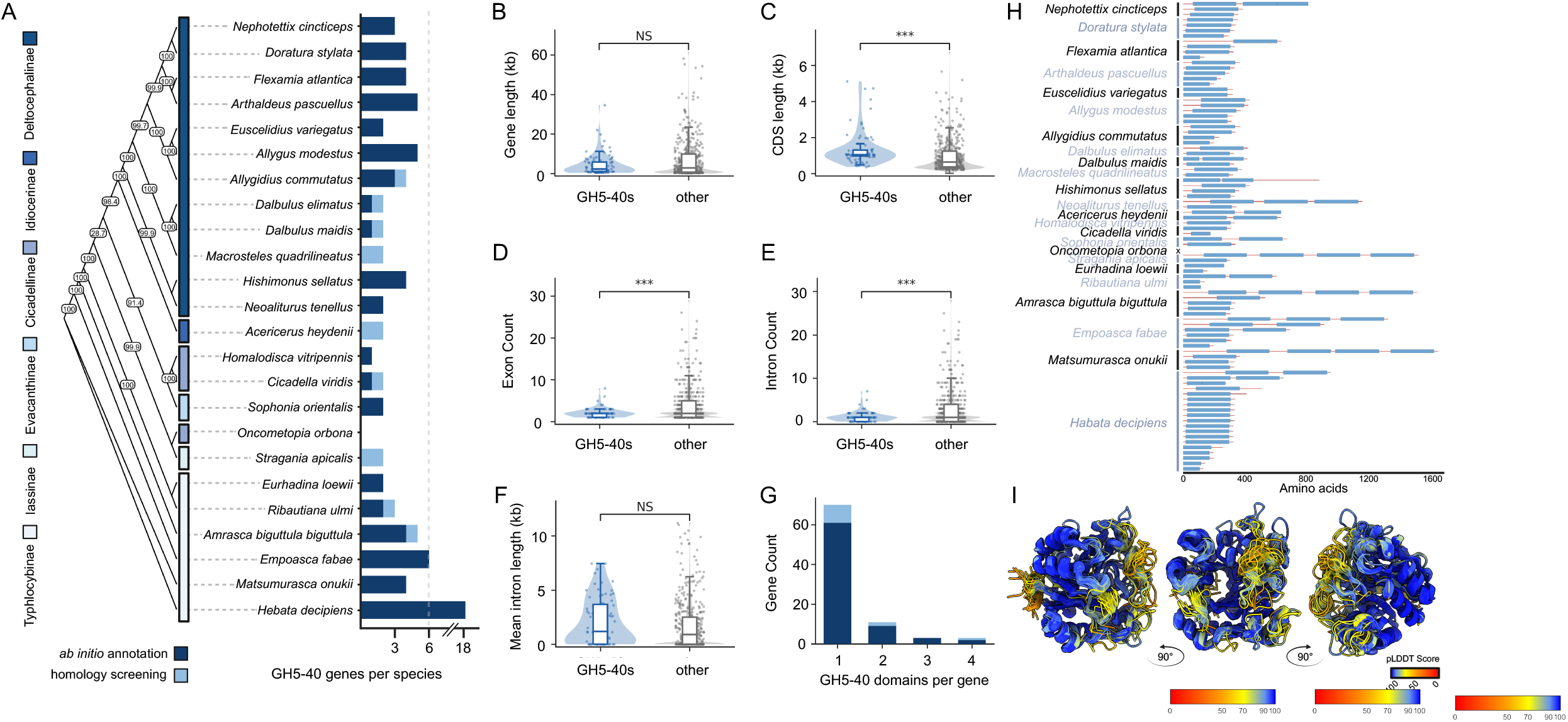
Distribution and expansion of GH5-40 across Cicadellidae. (A) Species tree of 24 leafhopper genomes with subfamily assignment and GH5-40 genes per species, split by recovery method (dark, *ab initio*; light, homology screening). Species tree inferred by maximum likelihood from 63 single-copy orthologs (21,520 amino-acid sites), node values are SH-aLRT/ultrafast bootstrap (%) from 1000 replicates. *(B-F) Architecture of GH5-40 loci (*n *= 87) versus all other annotated genes (*n = 1,039,190): gene length (B), CDS length (C), exon count (D), intron count (E), mean intron length (F). Boxes show median and interquartile range over violins of the full distribution. Two-sided Mann-Whitney U; NS, not significant; ***p < 0.001. (G) GH5-40 domains per gene, colored by identification method as in A. (H) Domain architecture of every GH5-40 protein, grouped by species and scaled by amino acid length. ‘x’ marks the absence of a recovered GH5-40 (Oncometopia orbona). (I) Structural superposition of all 113 predicted leafhopper GH5-40 domains, shown in three orientations. Color represents per-residue pLDDT confidence scores (blue, high; red, low)

Relative to all other annotated genes, GH5-40s were encoded by longer coding sequences (median 1,038 vs 636 bp) but were restricted to fewer exons and introns (**Fig. 1B-E**). Total gene length and mean intron length were indistinguishable from the background (**Fig. 1B,F**). Given the very large genomic background surveyed (*n* = 1,039,190 genes), we interpret this as a shift in distribution rather than relying on significance alone. Notably, unlike their bacterial homologs, leafhopper GH5-40 loci retained introns (**Fig. 1E,F**) and are transcribed in hosts, arguing against assembly contamination and indicating genuine integration while retaining compact, bacteria-like coding architectures (15).

Expansion of this subfamily appears to have occurred not only through increasing gene copy numbers but also through repeated copies of domains within genes. Seventeen genes encoded between 2 and 4 GH5-40 domains, including 11 two-domain, 3 three-domain, and 3 four-domain proteins (**Fig. 1G-I**). In total, multidomain genes were identified in 12 species spanning five subfamilies, with four-domain proteins present in distantly related Typhlocybinae (*Matsumurasca onukii* and *Amrasca biguttula biguttula*) and Iassinae (*Stragania apicalis*). This repeated occurrence of complete, multidomain architectures across deeply diverged lineages suggests that domain expansion has evolved multiple times independently following an initial acquisition (16), and has not been observed so extensively in other insect lineages to our knowledge.

To distinguish a single acquisition from multiple independent transfers, we reconstructed a phylogeny of all 113 catalytic domains together with 313 bacterial GH5-40 references from CAZy. Maximum-likelihood analysis placed all leafhopper domains within a single, strongly supported clade nested within Actinobacteria (**Fig. 2A**), consistent with descent from one horizontal acquisition retained through subsequent speciation (17, 18). Although domains from the same multidomain gene did not consistently resolve as sister lineages, the overall phylogeny robustly supports a single ancestral origin of the GH5-40 family, while the precise number and timing of internal domain-duplication events remain unresolved.

**Figure 2.**
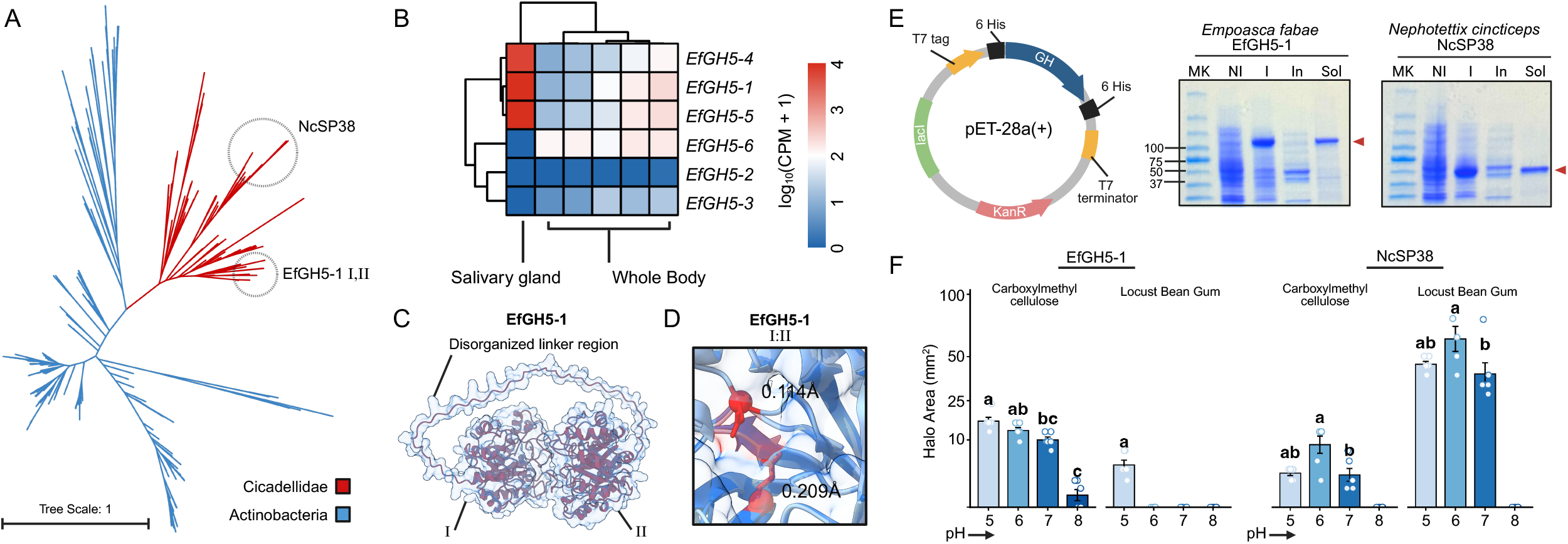
Origin, expression, structure, and activity of leafhopper GH5-40 enzymes. (A) Unrooted maximum-likelihood phylogeny of 113 leafhopper GH5-40 catalytic domains with 313 bacterial CAZy references (Q.pfam+R7 model; 1,000 ultrafast bootstraps; 447 aa alignment). Cicadellidae (red) form a single clade nested within Actinobacteria (blue). Clades of the EfGH5-1 domains (I, II) and NcSP38 are circled. (B) Expression of six *E. fabae* GH5-40 genes across one salivary-gland and five whole-body libraries, log_10_ (CPM + 1), with hierarchical clustering. (C) AlphaFold3 model of EfGH5-1 showing catalytic domains I and II joined by a disordered linker. (D) Superimposition of domains I and II at the catalytic cleft. Values are distances between paired catalytic residues (Å). (E) pET-28a(+) construct with N-terminal 6xHis-T7 tag and SDS-PAGE of recombinant EfGH5-1 and NcSP38 (MK, marker; NI, non-induced; I, induced; In, insoluble; Sol, soluble). Red arrows mark the target protein. (F) Halo areas on carboxymethylcellulose and locust bean gum at pH 5-8. Bars are mean ± SE (*n* = 6). Letters denote significant groups (one-way ANOVA, Tukey; α = 0.05).

We next asked whether multidomain GH5-40 enzymes are secreted and catalytically active. In *Nephotettix cincticeps*, the salivary protein NcSP38, previously identified by salivary proteomics, corresponds to the first catalytic domain of a full-length three-domain GH5-40 gene identified here (NcGH5-3; domain I; 99.3% identity, 288/290 residues). This finding reveals that NcSP38 is not a standalone enzyme, but rather part of a previously unrecognized multidomain protein. Consistent with a secretory role, three GH5-40 genes in the distantly related leafhopper *Empoasca fabae* were highly expressed in the only available salivary gland transcriptome (**Fig. 2B**). For one of these, EfGH5-1, structural modelling resolved two complete catalytic domains joined by a disordered linker (**Fig. 2C**). The two predicted catalytic glutamate pairs superimposed closely (0.114 and 0.209 Å; **Fig. 2D**), indicating that both domains present the conserved GH5 catalytic geometry.

Recombinant EfGH5-1 and the NcSP38 domain (NcGH5-3 domain I) were expressed, purified (**Fig. 2E**) and screened on polysaccharide plates. Despite the subfamily’s annotation as endo-β-1,4-mannanases (19), both hydrolyzed β-mannan and β-glucan but with contrasting preferences (**Fig. 2F**). EfGH5-1 favored carboxymethylcellulose (16.49 ± 1.50 mm^2^ at pH 5; mean ± SE, *n* = 6) over locust bean gum (4.19 ± 0.94 mm^2^), whereas NcSP38 favored locust bean gum (64.14 ± 9.68 mm^2^ at pH 6), exceeding its carboxymethylcellulose activity 6.8-fold. The two enzymes differed significantly on both substrates at multiple pH values (**Dataset S3**). As this qualitative diffusion screen samples only two enzymes, we interpret the contrast as an initial indication that salivary GH5-40 substrate preference may diversify across Cicadellidae, rather than as a family-wide demonstration.

## CONCLUSION

While retaining signatures of bacterial ancestry, leafhopper GH5-40 genes have undergone repeated gene expansion and independent elaboration of multidomain architectures. Although the subfamily has reported endo-β-1,4-mannanase activity, our data suggest that substrate preference may have broadened to other β-1,4-linked polysaccharides in at least some lineages. These enzymes illustrate how a single horizontal acquisition can become integrated into an animal genome and continue to diversify across speciation.

## MATERIALS AND METHODS

Briefly, 24 Cicadellidae assemblies (**Dataset S1**) were repeat-masked and annotated with BRAKER3 under a uniform protein-evidence set. GH5-40 domains were assigned by dbCAN HMM. Loci missed by *ab initio* prediction were recovered by miniprot homology search (*Dataset S2*). Catalytic domains were extracted, aligned with MAFFT, trimmed, and used for maximum-likelihood inference in IQ-TREE2 with 1,000 ultrafast bootstraps alongside all CAZy GH5-40 references. Architecture metrics were compared against each assembly’s longest-isoform gene set. EfGH5-1 and NcSP38 were modelled with AlphaFold3, expressed in *E. coli* from pET-28a(+), purified by Ni-NTA, and assayed on 0.2% carboxymethylcellulose and locust bean gum agar at pH 5-8 (**Dataset S3**).

### Genome assemblies and *ab initio* annotations

Twenty-four publicly available Cicadellidae genome assemblies spanning six subfamilies (Deltocephalinae, Idiocerinae, Cicadellinae, Evacanthinae, Iassinae, Typhlocybinae) were annotated. Assembly identifiers, NCBI accessions, and subfamily assignment for each genome are listed in Dataset S1. To ensure that gene models were standardized and directly comparable across species, every assembly was re-annotated under an identical workflow rather than using the annotations distributed with each assembly.

For each assembly, a species-specific *de novo* repeat library was generated with RepeatModeler v2.0.7 (20). The assembly was then soft masked with RepeatMasker v4.1.8 (21) using that library. Repeat libraries were built independently per genome, and no cross-species library was applied, to avoid lineage-specific repeat content biasing masking in any one taxon.

Masked assemblies were then annotated with BRAKER3 v3.0.8 (22). For comparative annotation across species, BRAKER3 was run under a uniform evidence set that excluded RNA-seq data so that gene prediction would not be biased by uneven transcriptomic coverage across taxa. Prediction inputs relied on a shared protein reference set (a custom 2.2-Gb Arthropoda proteome) supplied to GeneMark-ETP v1.0 (23) and GeneMark-EP+ v4.71 (24) for protein-guided training, with AUGUSTUS v3.5.0 generating final models and TSEBRA v1.1.1 (25) selecting transcripts. Protein alignment used DIAMOND v2.1.8 (26). Transcript model refinement used StringTie2 v2.2.1 (27). Annotation completeness was assessed against the hemiptera_odb10 lineage with BUSCO v5.4.5 (28), Miniprot v0.12 (29), and Compleasm v0.2.6 (30) implemented within BRAKER3. Only the longest isoform per locus was retained for all downstream analyses.

### Species tree inference

Orthogroups were inferred across the 24 annotated proteomes with OrthoFinder v3.1.0 (31). Single-copy orthologues shared across taxa were aligned with MAFFT v7.533 (32), trimmed with trimAl v1.4 (33), concatenated, and analyzed in IQ-TREE2 v3.0.1 (34) with the best-fit substitution model selected by ModelFinder (35) and node support from 1,000 ultrafast bootstrap replicates (36). The resulting topology in relation to previously established subfamilies is shown in Fig. 1A.

### GH5-40 identification and domain annotation

Carbohydrate-active enzyme domains were annotated across all 24 proteomes with dbCAN (37), and proteins carrying at least one GH5 subfamily 40 (GH5-40) HMM assignment were retained. For every retained protein, dbCAN3 HMM coordinates defined catalytic domain boundaries. Proteins carrying more than one non-overlapping GH5-40 were classified as multidomain (Dataset S2). Two additional fields were then assigned per gene: (1) domain completeness, which distinguishes domains spanning the full HMM (complete) from truncated ones (partial), and (2) gene status, which distinguishes structurally intact models (intact) from those carrying in-frame stops or frameshifts (pseudogenes).

### Homology-based recovery of unannotated loci

To distinguish any genuine absence of GH5-40 from missing *ab initio* predictions, every assembly was additionally screened by protein alignment with Miniprot v0.12, using a curated HMM profile of the previously recovered GH5-40 proteins. Identified candidate loci not overlapping an existing BRAKER3 model were then extracted and reviewed with dbCAN. Loci returning a GH5-40 assignment were retained. This recovered 12 additional genes across nine species, which were assessed with the 75 *ab initio* models in all downstream analyses. These two gene sets are distinguished by the annotation method in Dataset S2 and by color in Fig. 1A and 1G. Applying the identical screen to *Oncometopia orbona* returned no additional GH5-40 locus, and this genome is reported to be absent of GH5-40.

### Gene architecture analysis

For each of the 24 assemblies, gene-level architecture metrics were extracted from the BRAKER3 annotation for all longest-isoform models, including the total gene length (including introns), coding sequence (CDS) length, exon count, intron count, and mean intron length. Coordinates were parsed with the rtracklayer (38) and GenomicFeatures (39) packages in R (40). GH5-40 loci were compared against all annotated genes (background *n* = 1,039,190) using two-sided Mann-Whitney U tests.

### GH5-40 domain phylogeny

All GH5-40 protein sequences were retrieved from the CAZy database. Leafhopper catalytic domains were excised at the defined domain boundaries denoted by dbCAN3 and treated as independent sequences, so that multidomain proteins with inter-domain linkers did not distort the alignment. Leafhopper domains (*n* = 113) and CAZy reference domains (*n* = 313) were aligned with MAFFT v7.533 under the L-INS-i algorithm, trimmed with trimAl v1.4, and analyzed by maximum likelihood in IQ-TREE2 v3.0.1 with the best-fit substitution model selected by ModelFinder. Node support was estimated from 1,000 ultrafast bootstrap replicates.

### Tissue-specific expression

Expression of the six *Empoasca fabae* GH5-40 genes was assessed from publicly available RNA-seq datasets. The salivary gland dataset (SRA accession SRR275587) (41) and five whole-body datasets (SRR17845030–SRR17845034) (42) were adapter and quality-trimmed with fastp v1.0 (43) and mapped to the *E. fabae* reference assembly in paired-end mode with HISAT2 v2.2.1 (44). Alignments were coordinate-sorted and indexed with SAMtools v1.17 (45), and gene-level counts generated with HTSeq v2.0.2 (46). Counts were normalized to counts per million (CPM), log_10_ (CPM + 1)-transformed, and hierarchically clustered by row and column.

### Structural modeling

Three-dimensional structures of EfGH5-1 and NcSP38 were predicted with AlphaFold3 (47). Model quality was assessed and the top ranked prediction were retained. Structural superimposition of the two EfGH5-1 catalytic domains and active site geometry were computed and visualized in UCSF ChimeraX v1.11.1 (48). The disordered linker joining the two domains was delimited by the region between the two dbCAN3 GH5-40 identified domains and validated by the presence of a low pLDDT score.

### Gene synthesis, cloning, and heterologous expression

Coding sequences had signal peptides removed and were then codon-optimized for *Escherichia coli*, synthesized, and cloned into pET-28a(+) with an N-terminal 6xHis tag (Twist Bioscience, San Francisco, CA, USA). The two enzymes assayed in this report were the *E. fabae* 2-domain GH5-40 (EfGH5-1) and the *N. cincticeps* single domain GH5-40 (NcSP38) (the deposited single domain protein BAQ94505.1, which corresponds to the first catalytic domain of the full-length 3-domain gene recovered here at 99.3% identity). Constructs were then transformed into *E. coli* LEMO21(DE3) (New England Biolabs, Ipswich, MA, USA) by heat shock and selected on LB agar with kanamycin (50 µg/mL). Single colonies were grown in LB with kanamycin (50 µg/mL) and chloramphenicol (34 µg/mL) at 37 °C, 220 rpm, to mid-log phase (OD_600_ = 0.6). Expression was induced with 0.5 mM IPTG followed by 18 h of shaking at 16 °C. Cells were harvested, resuspended in lysis buffer (50 mM NaH_2_PO_4_, 300 mM NaCl, 10 mM imidazole, pH 8.0), and disrupted by sonication (6 x 30 s on / 30 s off at 35% amplitude; Fisherbrand Model 505 Sonic Dismembrator, Thermo Fisher Scientific, Waltham, MA, USA). Soluble fractions were recovered by centrifuge (12,000 x g, 30 min, 4 °C) and purified over Ni-NTA agarose (QIAGEN, Hilden, Germany) with elution across a gradient concentration of 50-250 mM imidazole. Protein was quantified using a Quick Start Bradford assay (Bio-Rad, Hercules, CA, USA).

Final purity was assessed by SDS-PAGE on 4-15% Mini-PROTEAN TGX precast gels (Bio-Rad) stained with Bio-Safe Coomassie (Bio-Rad). For western blotting, proteins were transferred to 0.2-µm PVDF membranes (70 V, 400 mA, 2 h), blocked in 5% milk, and probed with anti-His_6_-HRP conjugate (Roche, Basel, Switzerland; Ref 11965085001) at 1:500 in 3% BSA. Signal was developed with Clarity Max Western ECL substrate (Bio-Rad) and imaged on an Azure c300 (Azure Biosystems, Dublin, CA, USA). Ponceau S served as a loading control.

### Polysaccharide activity assays

Endo-β-1,4-glucanase and endo-β-1,4-mannanase activities were assayed on agar plates. Substrate plates contained either carboxymethylcellulose sodium salt (CMC; 0.2% w/v) or locust bean gum (LBG; 0.2% w/v) (both Thermo Fisher Scientific) in 1.5% w/v agar buffered to pHs 5, 6, 7, or 8 (50 mM sodium citrate for pH 5 and 6; 50 mM sodium phosphate for pH 7 and 8). Equal 10 ul droplets (concentration 2.5 µM /ul) of purified protein were spotted onto agar surfaces. Plates were incubated at 28 °C for 24 h, then removed and stained with 0.1% w/v Congo red for 15 min. After staining, plates were then destained with 1 M NaCl and neutralized with 1% v/v HCl. Cleared halos were measured and halo area (mm^2^) was quantified. Six replicate halos were measured per enzyme, substrate, and pH combination (*n* = 6). All measurements are provided in Dataset S3.

## DATA, MATERIALS, AND SOFTWARE AVAILABILITY

Assembly accessions, GH5-40 protein sequences, and substrate assay data are provided in Datasets S1-S3. Extended methods and citations are available in the SI Appendix.

## AUTHOR CONTRIBUTIONS

Conceptualization, J.M. and E. P.-L.; methodology, J.M. and E.I.F; investigation, J.M., A.A.S., and E.P.-L.; data curation and formal analysis, J.M.; visualization, J.M.; resources and funding acquisition, E.P.-L.; project administration and supervision, E.P.-L.; writing original draft, J.M.; All authors contributed to review & editing of this manuscript.

## ACKNOWLEDGMENTS

We are thankful to resources provided by Université Laval’s high-performance computing infrastructure at the Institute of Biology Intégrative et des Systèmes (IBIS). Figures were created with BioRender.com. J.M. thanks NSERC for the PhD scholarship.

## SUPPLEMENTARY DATASETS S1 TO S3

### Dataset S1 (separate file). Cicadellidae genome assemblies analyzed in this study

Information for each assembly includes the species, subfamily, assembly identifier, NCBI assembly accession, and the number of GH5-40 genes, GH5-40 catalytic domains, genes recovered by *ab initio* annotation, genes recovered by

### homology screening, and multidomain genes

Dataset S2 (separate file). List of leafhopper GH5-40 genes. Information for each gene includes the species, subfamily, assembly identifier, gene identifier, genomic location (contig with position start/end and strand +/-), GH5-40 domain count, protein length, dbCAN3 domain coordinates, annotation method (*ab initio* or miniprot homology screening), gene and domain completeness, and the full protein sequence.

### Dataset S3 (separate file). Polysaccharide hydrolysis assay data for EfGH5-1 and NcSP38

Sheet 1 includes the individual halo measurements for each enzyme across substrates and pH and Sheet 2 includes a summary of means, standard deviations, and standard errors of each halo area.

